## Supplementary material for "A multi-hierarchical approach reveals D-serine as a hidden substrate of sodium-coupled monocarboxylate transporters": Figure supplements

**Figure supplement 1. Wiriyasermkul, et al.**

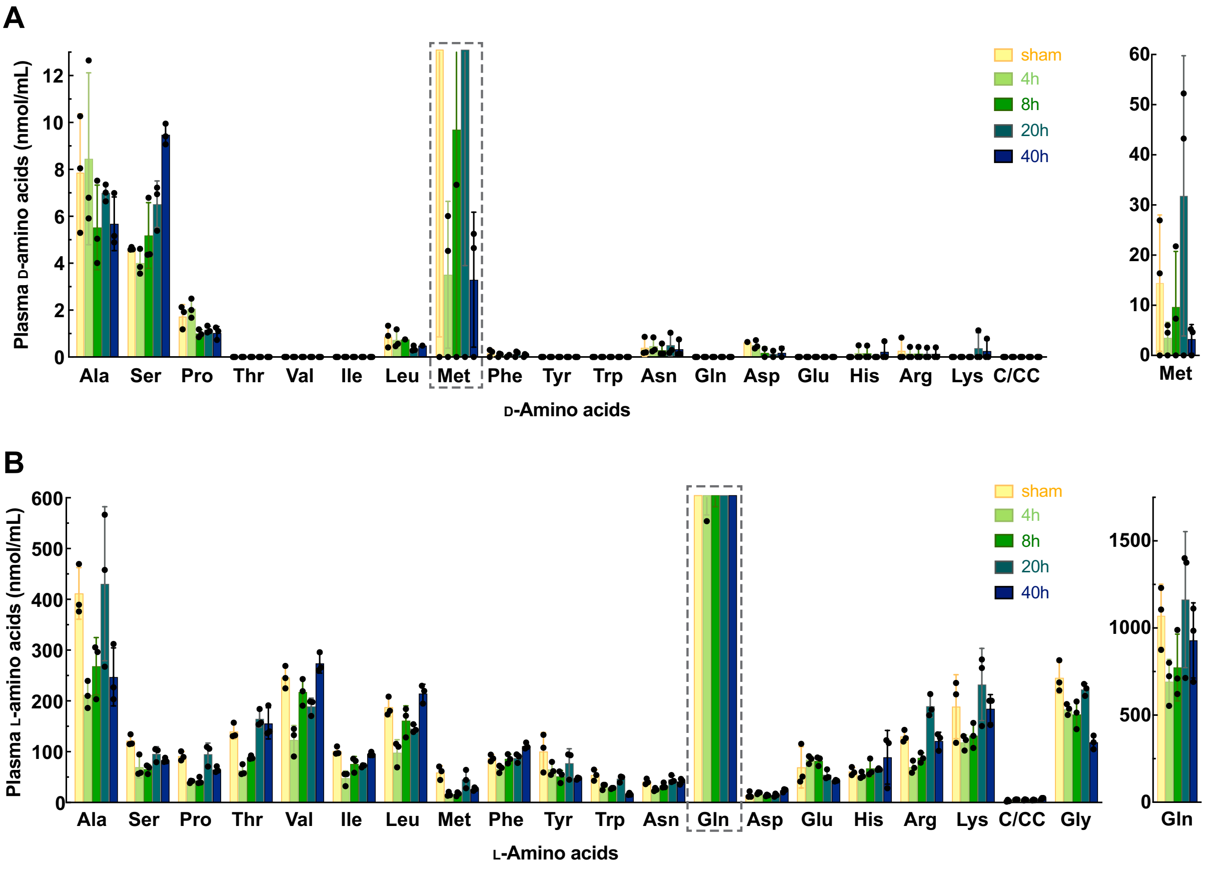


**Figure supplement 1. Enantiomeric profiles of D- and L-amino acids in plasma of the IRI model**

Plasma samples were collected from the mice after ischemia operation for 0, 4, 8, 20 and 40 hours or sham operation. Both D- and L- enantiomers of twenty amino acids were measured by 2D-HPLC and plotted as mean ± SD. (**A**) Concentration of plasma D-amino acids. Graphs of D-Met are shown separately for the proper resolution. (**B**) Concentration of plasma L-amino acids. Graphs of D-Gln are shown separately for the proper resolution. Glycine plots are included in **B**. C/CC: cysteine or cystine. n = 3.

### Figure supplement 2. Wiriyasermkul, et al.

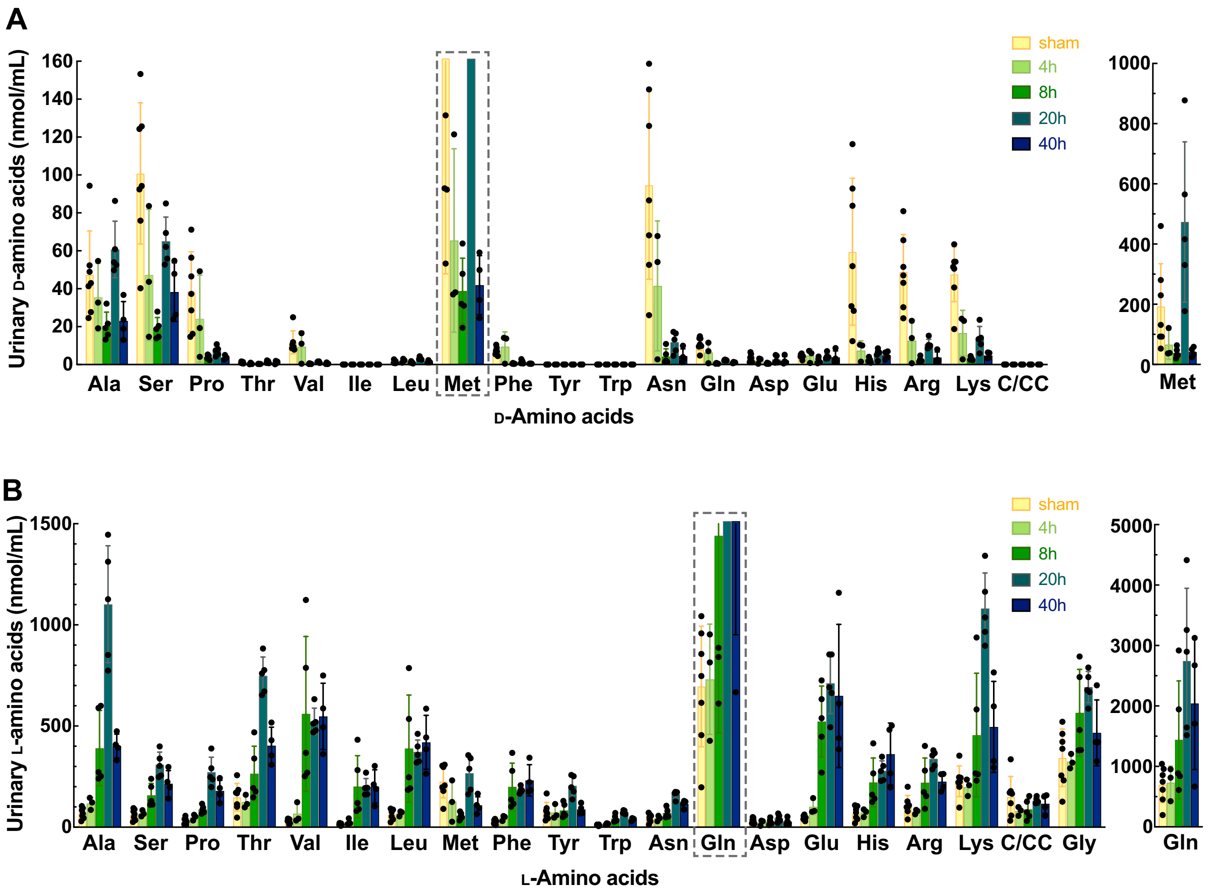


**Figure supplement 2. Enantiomeric profiles of D- and L-amino acids in urine of the IRI model**

Urine samples were collected from the mice after ischemia operation for 0, 4, 8, 20 and 40 hours or sham operation. Both D- and L- enantiomers of twenty amino acids and were measured by 2D-HPLC and plotted as mean ± SD. (**A**) Concentration of urinary D-amino acids. Graphs of D-Met are shown separately for the proper resolution. (**B**) Concentration of urinary L-amino acids. Graphs of D-Gln are shown separately for the proper resolution. Glycine plots are included in **B**. C/CC: cysteine or cystine. n = 3 – 7.

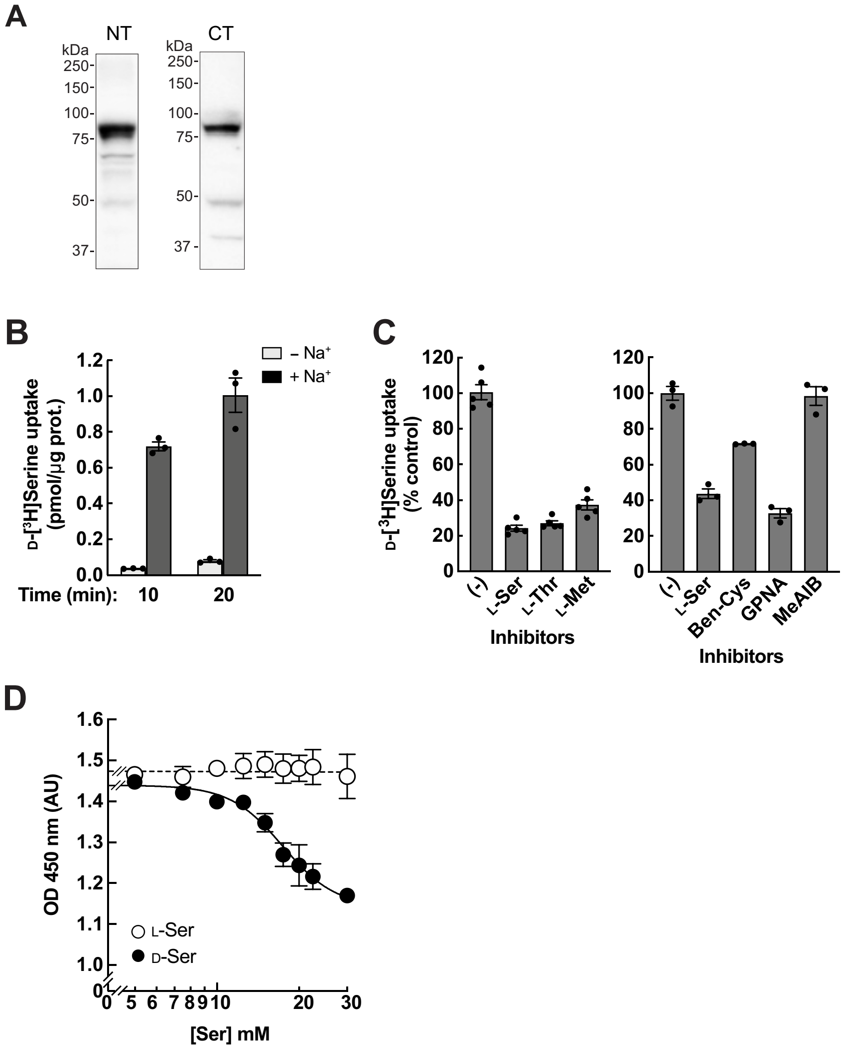


**Figure supplement 3. Characterization of ASCT2 as a D-serine transporter in HEK293 cells**

(**A**) Western blot of Asct2 from renal BBMVs of the normal mice. Asct2 was detected by anti-Asct2(NT) (left) or anti-Asct2(CT) (right) antibodies. (**B**) Transport of 10 μM D-[$^3$H]serine was measured for 10 or 20 min in HEK293 cells in the presence or absence of $Na^+$. Bar graph = mean ± SEM; n = 3. (**C**) Inhibition of D-[$^3$H]serine transport by several compounds. Left: 5 μM D-[$^3$H]serine uptake was measured for 10 min in the presence or absence of 2 mM L-amino acids. Bar graph = mean ± SEM; n = 5. Right: 2 μM D-[$^3$H]serine uptake was measured in the presence or absence of 1 mM inhibitors; Ben-Cys: *S*-benzyl-L-cysteine, a non-specific inhibitor of ASCT2; GPNA: L-γ-glutamyl-*p*-nitroanilide, an inhibitor of ASCT2, SNATs and LATs; MeAIB: 2-(methylamino)isobutyric acid, a system A inhibitor. Bar graph = mean ± SEM; n = 3. (**D**) Cell-growth measurement (XTT assay) of HEK293 cells treated with either L-serine or D-serine 5 – 30 mM for two days. D-Serine inhibition curve was fitted to non-linear regression of $\log_{10}$[D-Ser] v.s. cell growth, resulting in $IC_{50}$ of 17.4 ± 1.05 mM and minimum growth at OD 450 nm of 1.15 arbitrary units (AU). Dot plot = mean ± SEM; n = 3 – 5.

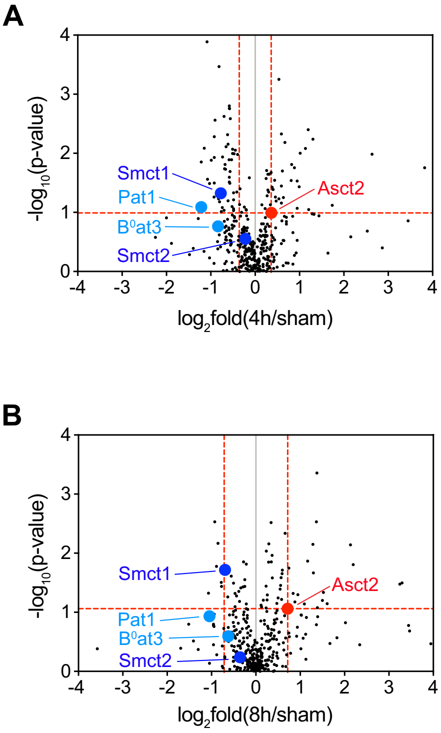


**Figure supplement 4. Expressions of Smct1, Smct2, Pat1, and $B^0$at3 in the volcano plots of membrane transport proteins**

Volcano plot of 389 membrane transport proteins as identically shown in Fig. 4A. Positions of Smct1, Smct2, Pat1, and $B^0$at3 are indicated. (**A**) The plot of proteins from 4h/sham. (**B**) The plot of proteins from 8h/sham.

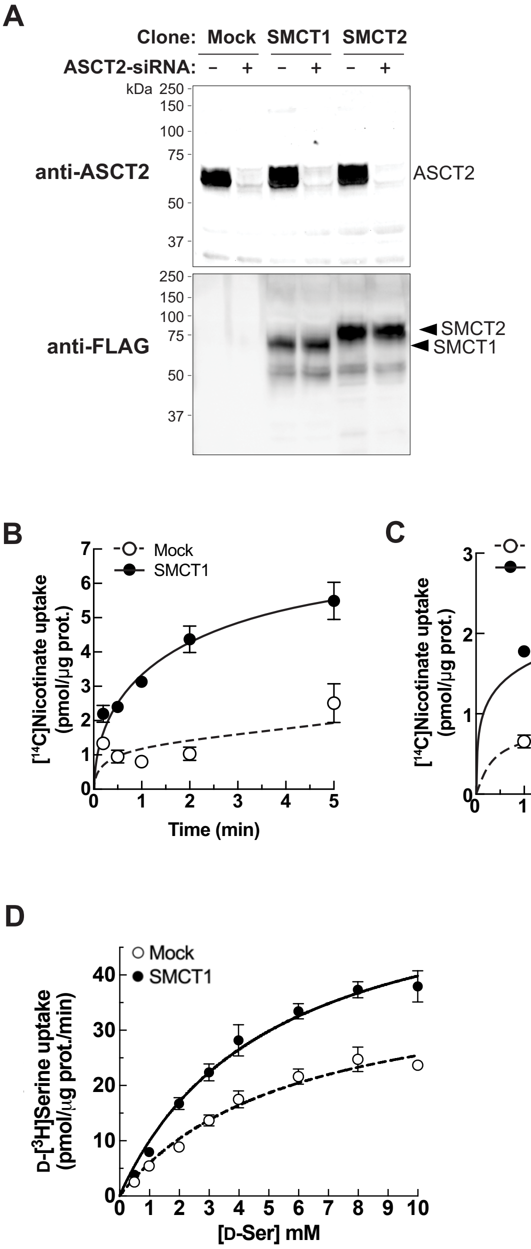


**Figure supplement 5. SMCT1 and SMCT2 functions in FlpInTR-SMCT1 and FlpInTR-SMCT2 stable cell lines**

(**A**) Western blot of membrane fractions from FlpInTR-Mock, -SMCT1, and -SMCT2 cells with ASCT2-siRNA transfection. Flp-In T-REx 293 stably expressing SMCT1 (SMCT1) and

SMCT2 (SMCT2), as well as Mock, were transfected with ASCT2-siRNA for two days. Membrane proteins were extracted from crude membrane fractions and subjected to Western blot analysis. ASCT2 knockdown efficiency was evaluated by anti-ASCT2 antibody. Expression of SMCT1 and SMCT2 were verified by anti-FLAG antibody. (**B**) – (**C**) Evaluation of SMCT1 and SMCT2 functions by transport assay. Time course of 50 μM [$^{14}$C]nicotinate uptake was measured for 0.5 – 5 min in FlpInTR-SMCT1 (**B**) and FlpInTR-SMCT2 (**C**) cells, compared to Mock cells. Dot plot = mean ± SEM; n = 3. (**D**) Raw data of Fig. 5E: D-[$^{3}$H]serine uptake in FlpIn293TR-SMCT1 compared to FlpIn293TR-Mock cells. The uptake values were fitted to Michaelis-Menten plot. Dot plot = mean ± SEM; n = 3 – 4.

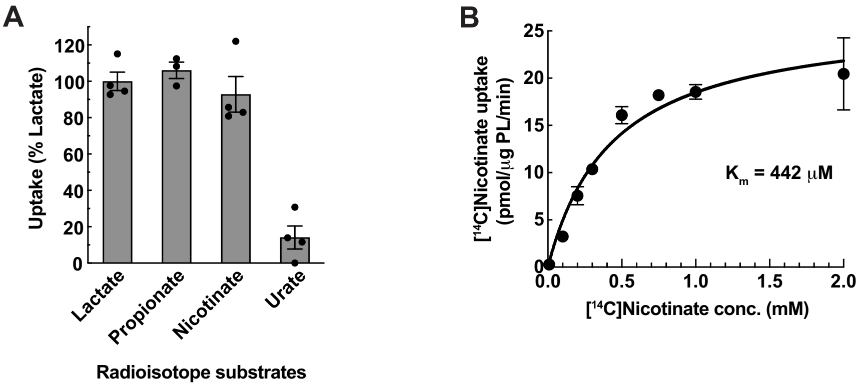


**Figure supplement 6. SMCT1 function in SMCT1-reconstituted proteoliposomes**

(**A**) Transport of SMCT1 canonical substrates in SMCT1-reconstituted proteoliposome (SMCT1-PL). Uptake of 50 μM radiolabeled substrates was measured for 5 min in $Na^+$-containing buffer and subtracted with the uptake in $Na^+$-free buffer. Urate is used as a negative control. Bar graph = mean ± SEM; n = 3 – 4. (**B**) Concentration dependence of [$^{14}$C]nicotinate uptake in SMCT1-PL. Uptake of [$^{14}$C]nicotinate (0.01 – 2 mM) by SMCT1-PL was measured for 3 min in $Na^+$-containing buffer and then subtracted with the uptake in $Na^+$-free buffer. The graph was fitted to Michaelis-Menten plot with the apparent $K_m$ of 442 ± 94 μM and $V_{max}$ of 26.7 ± 2.23 pmol/μg PL/min. Dot plot = mean ± SEM; n = 3 – 4.

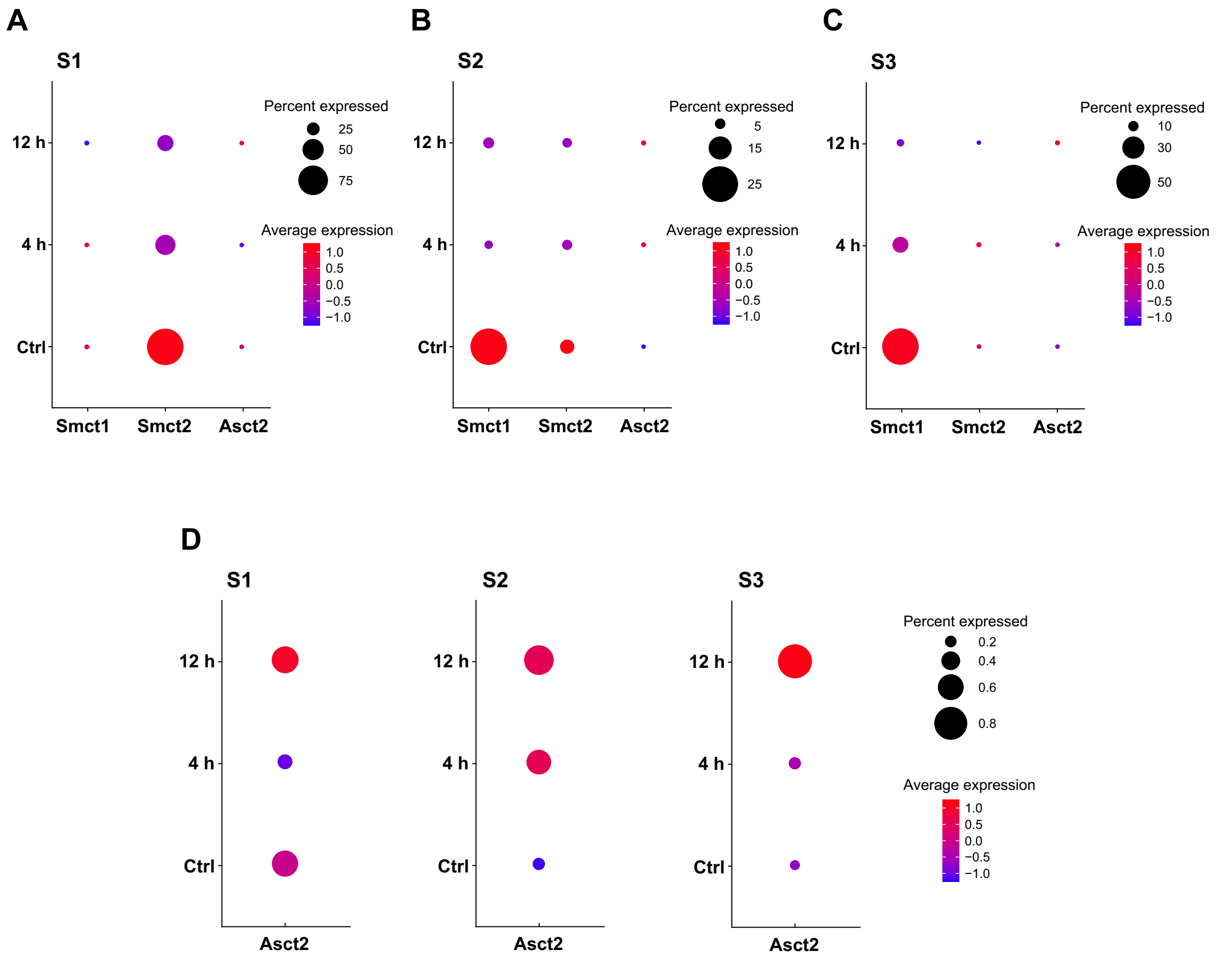


**Figure supplement 7. Expression of Smct1, Smct2 and Asct2 in renal proximal tubules of the IRI model by snRNA-seq**

Data of Smct1, Smct2 and Asct2 expression in the IRI model by snRNA-seq were obtained from the open-sourced dataset. Bubble plots indicate the expressions of Smct1, Smct2 and Asct2 in clusters of proximal tubule segments 1 (S1) (**A**), 2 (S2) (**B**), and 3 (S3) (**C**). The mRNA expressions at early IRI stages (4 and 12 hours) were compared to the sham controls (Ctrl). (**D**) The zoom-in plots of Asct2 from (**A**) – (**C**). "Percent Expressed" represents the percentage of cells expressing each gene in each cluster. "Average Expression" indicates the scaled expression level: 0, the mean expression of a gene across cells; the positive variance, increased expression; the negative variance, decreased expression.

### Figure supplement 8. Wiriyasermkul, et al.

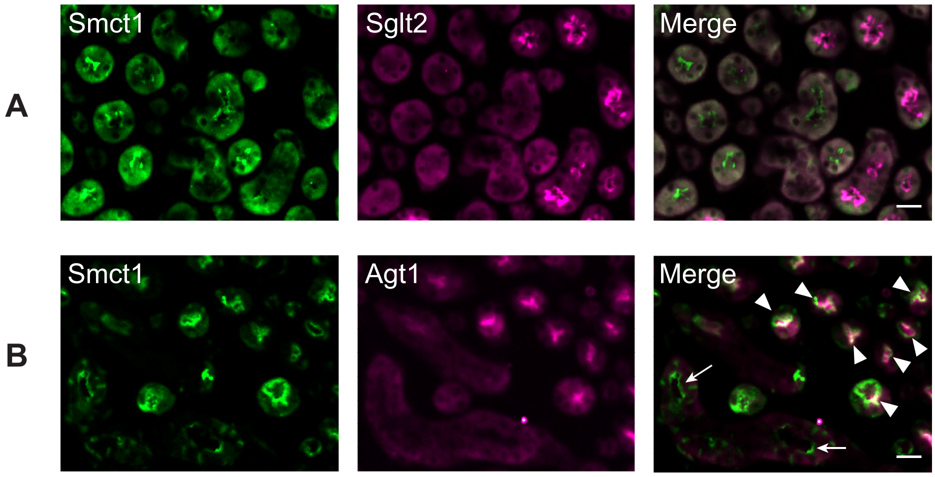


**Figure supplement 8. SMCT1 is mainly localized at the apical membrane of renal proximal tubular S3 segment**

Localization of Smct1 in the mouse kidney was determined by immunofluorescence staining. Mouse kidney slide was co-stained with anti-Smct1 (Smct1; green) antibody and protein markers for renal proximal tubule segments: anti-Sglt2 antibody (**A**: Sglt2, apical membrane marker of S1 + S2 segments) or anti-Agt1 antibody (**B**: Agt1, apical membrane marker of S3 segment). Merge images are shown in the right panel. Arrowheads indicate co-localization of the proteins. Arrow shows some parts of the faint Smct1 without Agt1 co-localization. Scale bar = 20 µm.

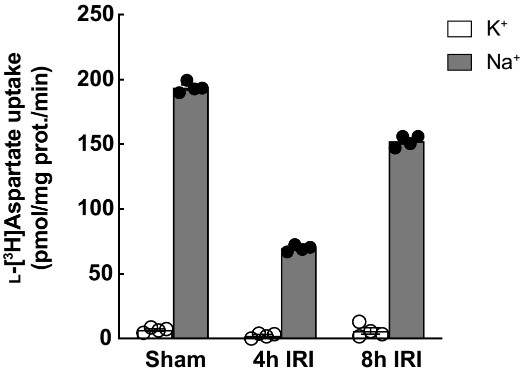


**Figure supplement 9. Functional L-aspartate transport in BBMVs from IRI model**

Function of L-aspartate transport, chiefly by excitatory amino acid transporter Slc1a1/Eaac1, in the IRI model was determined by L-[$^3$H]aspartate transport. Uptake of 10 μM L-[$^3$H]aspartate in renal BBMVs isolated from IRI model was measured for 1 min in $Na^+$-free buffer ($K^+$) or $Na^+$-containing buffer ($Na^+$). Bar graph = mean ± SEM; n = 4.

**Table supplements 1-3 are provided separately in the Excel files.**

Table supplement 1: Proteomics of BBMVs from the IRI model

Table supplement 2: Annotation of membrane transport proteins from BBMV proteomics

Table supplement 3: Proteomics of HEK293 membrane
